# Distinct Brain Mechanisms for Conflict Adaptation Within and Across Conflict Types

**DOI:** 10.1101/2021.05.30.446264

**Authors:** Guochun Yang, Kai Wang, Weizhi Nan, Qi Li, Ya Zheng, Haiyan Wu, Xun Liu

## Abstract

Cognitive conflict, like other cognitive processes, shows the characteristic of adaptation, i.e., conflict effects are attenuated when immediately following a conflicting event, a phenomenon known as the conflict adaptation effect (CAE). One important aspect of CAE is its sensitivity to the intertrial coherence of conflict type, i.e., behavioral CAE occurs only if consecutive trials are of the same conflict type. Although reliably observed behaviorally, the neural mechanisms underlying such a phenomenon remains elusive. With a paradigm combining the classic Simon task and Stroop task, this fMRI study examined neural correlates of conflict adaptation both within and across conflict types. The results revealed that when the conflict type repeated (but not when it alternated), the CAE-like neural activations were observed in dorsal anterior cingulate cortex, inferior frontal gyrus, superior parietal lobe, etc. (i.e., regions within typical task-positive networks). In contrast, when the conflict type alternated (but not when it repeated), we found CAE-like neural deactivations in a range of regions including bilateral superior and medial frontal gyri, bilateral angular cortex, bilateral temporal cortices, etc. (i.e., regions within the typical task-negative network). Moreover, this CAE-like neural deactivation predicts behavior performance. Network analyses suggested that these regions (for CAE-like neural activities within and across conflict type[s] respectively) can be clustered into two antagonistic networks. This evidence suggests that our adaptation to cognitive conflicts within a conflict type and across different types may rely on these two distinct neural mechanisms.

## 1. Introduction

Adaptation is an important property of many cognitive and neural processes which can occur at different cognitive levels when we are repetitively exposed to the same type of stimuli (Clifford & Palmer, 2018; Thompson & Burr, 2009; Zaske, Schweinberger, Kaufmann, & Kawahara, 2009). At higher levels of cognition, adaptation has been used as a research tool to probe the process of cognitive control, typically via adaptations in conflict processing. The conflict effect decreases after encountering an incongruent event relative to encountering a congruent event, a phenomenon known as the conflict adaptation effect (CAE) (Duthoo, Abrahamse, Braem, Boehler, & Notebaert, 2014; Gratton, Coles, & Donchin, 1992). Importantly, behavioral CAEs are highly sensitive to the coherence of the conflict type in adjacent trials, i.e., CAEs happen only when consecutive trials belong to the same conflict type (e.g., a Stroop type of conflict vs. a Simon type of conflict). Although this sensitivity of the CAE has been extensively reported and discussed at behavioral level (for a review, see Braem, Abrahamse, Duthoo, & Notebaert, 2014), the corresponding neural mechanisms are still unclear.

A behavioral CAE is commonly defined as the reaction time (RT) difference between the conflict effect after a congruent trial and the conflict effect after an incongruent trial, as described by the following equation:

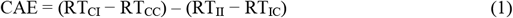

where C and I are the abbreviations of congruent and incongruent, respectively (Nieuwenhuis et al., 2006). To investigate sensitivity of the CAE to intertrial coherence on conflict type, the CAE-related brain activities in both within-type and across-type conditions need to be examined and compared (eight conditions). However, previous studies have examined brain areas showing a CAE-like neural activation mainly within the same conflict type (Carter et al., 2000; Chechko, Kellermann, Schneider, & Habel, 2014; Chun, Park, Kim, Kim, & Kim, 2017; Egner & Hirsch, 2005b), and the neural mechanisms understanding the loss of CAE in across-type conditions have rarely been examined. Therefore, in this study, to explore the full picture of the neural correlates of CAEs, especially the neural mechanisms underlying sensitivity to conflict type, it was necessary to perform the analysis based on its definition in both conflict type repetition and alternation conditions (see Methods for details).

To date, there have been only a limited number of event-related potential (ERP) studies and region of interest (ROI) studies attempting to reveal the mechanisms underlying the conflict-type sensitivity. N2 and P3, two components corresponding to the mental processing of conflict detection and attention allocation (Clayson & Larson, 2011), were found to show a CAE only when the consecutive conflict sequences were repeated (Q. Li et al., 2015; Z. Li et al., 2021). In addition, an ROI-based functional magnetic resonance imaging (fMRI) study observed the conflict type sensitivity functions that focused on the conflict detection region (i.e., anterior cingulate cortex [ACC]) and executive control regions (i.e., premotor cortex and dorsolateral prefrontal cortex [DLPFC]) (Kim, Chung, & Kim, 2010, 2012). These studies together implied that the lack of behavioral CAEs in conflict-alternation conditions might reflect the absence of conflict detection, attention allocation and executive control mechanisms in alternating conflict type sequences. However, the low spatial resolution of ERP technology (Q. Li et al., 2015; Z. Li et al., 2021) and the ROI-based method (Kim et al., 2010, 2012) cannot describe the whole picture of neural processing in CAEs sensitive to conflict types. It remains possible that other CAE related brain areas reported in previous studies, such as the superior parietal lobe (SPL) (Egner, Delano, & Hirsch, 2007) and inferior frontal gyrus (IFG) (Egner, 2011), may also show conflict type sensitive CAE.

The current study aimed to elucidate the neural mechanisms of the sensitivity of the CAE to conflict type with a whole-brain exploratory method. We adopted a Stroop-color-Simon paradigm and collected fMRI data during the task performance. This paradigm has been reported to be valid in producing robust behavioral and neural conflict-type sensitive CAEs (Liu, Park, Gu, & Fan, 2010; K. Wang, Li, Zheng, Wang, & Liu, 2014). We hypothesize that the conflict processing related brain areas, such as the cingulo-opercular and frontoparietal regions, would show CAE-like neural activities (mirroring behavioral CAEs) only in conflict type repetition but not in conflict type alternation conditions. Additionally, we predict that the brain regions showing conflict-type sensitive CAEs could predict the behavior.

## 2. Methods

### 2.1. Participants

Twenty right-handed volunteers (8 males and 12 females, average age: 21.7±1.6 years) took part in the experiment. The sample size was decided based on previous fMRI studies detecting similar CAE effects (Chun et al., 2017; Kim et al., 2012; Purmann & Pollmann, 2015). All participants were healthy, with normal or corrected-to-normal visual acuity and were free of psychiatric or neurological history. Before the experiment, all participants signed an informed consent form that was approved by the Institutional Review Board of the Institute of Psychology, Chinese Academy of Sciences. All participants were compensated for their participation. One participant was removed from the statistical analysis because of excessive head motion (rotation > 2 degrees in two runs).

### 2.2. Apparatus, Stimuli, and Procedure

The paradigm was adopted from previous studies (Liu et al., 2010; K. Wang et al., 2014) and modified for the fMRI experiment (see Figure 1). Stimulus presentation was controlled by E-Prime 2.0 (Psychological Software Tools, Inc., Pittsburgh, PA, USA). The stimulus was a center-displayed diamond (visual angle 4.9° × 4.9°) with half (a triangle) painted either red or blue. The triangle pointed in one of four directions (left, right, up, and down). A Chinese character indicating a color (i.e., “红” meaning red, or “蓝” meaning blue) or having a neutral meaning (i.e., “杯” means a cup, and “莫” means “do not”; these two words were selected because they had similar font structure with “红” and “蓝”, respectively) displayed in black ink, was overlaid in the center of the diamond. All stimuli were presented on a gray background. Before scanning, the participants were trained to become familiar with the task. The participants were allowed to enter the scanner to perform a formal test when their training accuracy reached 90%. Color-response mapping was counterbalanced across participants.

**Figure 1.**
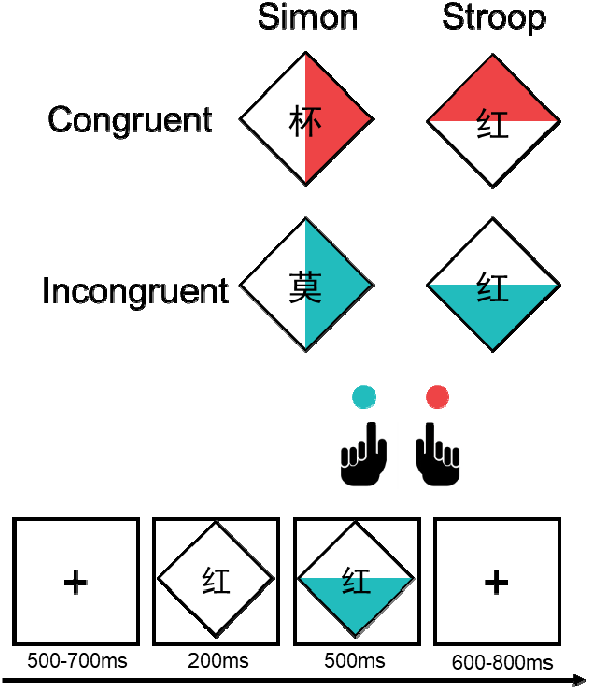
Experimental design and procedures. Participants were asked to respond to the color of the triangle and ignore any other information.

There were two types of conflicts during the test. In the Stroop conflict, the word was either “red” or “blue”, and the color of the triangle was either red or blue. Whether or not the character matched the color of the triangle formed the Stroop congruent (StC) and Stroop incongruent (StI) conditions, respectively. In addition, the triangle always pointed up or down to avoid a combination with a Simon conflict. In the Simon conflict, the colored triangle pointed left or right. The consistency between the orientation of the triangle and the response hand (left or right) determined the Simon congruency, i.e., Simon congruent (SmC) or Simon incongruent (SmI). In addition, the overlaying word was always color-irrelevant (e.g., “杯” meaning cup) to avoid a combination with a Stroop conflict. The participants were instructed to make a left or right key press based on the color of the stimulus (red or blue) while ignoring other information and to respond as quickly and accurately as possible. From the perspective of the participants, there were no differences between the Simon and Stroop tasks. Therefore, there was no task switching between different conflicts.

The participants performed four test sessions. Each session consisted of 162 trials listed in a pseudorandom fashion, with equal numbers of StI, StC, SmI and SmC trials intermixed randomly, and equal probability of each secondary trial sequence (e.g., StC-SmI, SmC-StC). The pseudorandom lists were generated with the AlphaSim function of AFNI software. Each trial lasted 2000 ms, with a prestimulus fixation icon presented centrally for 100∼300 ms, followed by a white diamond with a character in the middle (700 ms); then, the task stimulus (a colored triangle) appeared 200 ms after the onset of the diamond and lasted for 500 ms, after which a poststimulus fixation icon was presented for the remainder of the trial. The participants were allowed a maximum of 1500 ms from the onset of the target display to respond. In addition, to better estimate the event-related fMRI signals, 55 blank trials with only the fixation icon, each lasting 2000 ms, were inserted into each session, dividing each long run into multiple mini-blocks. The number of fixation trials between mini-blocks followed the exponential distribution.

### 2.3. Behavioral Data Analysis

Data were analyzed with dependent variables of both reaction time (RT) and error rate (ER). To avoid misleading potential conflicting RT and ER results, we also calculated the linear integrated speed-accuracy score (LISAS), an index that has been proven to efficiently account for the variance in behavioral measures (Vandierendonck, 2017). The LISAS was calculated with the following equation:

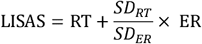

The first trial of each mini-block (10.5%), error trials (3.7%), correct trials after an error trial (3.2%), and trials with RTs beyond 3 standard deviations (SDs) of the mean or shorter than 200 ms (0.4%) were excluded before analyzing the interaction between the previous congruency and the current congruency (i.e., the CAE). We conducted three-way repeated-measures analyses of variance (ANOVAs) of consecutive conflict type (2, repetition vs. alternation) × previous congruency (2, congruent vs. incongruent) × current congruency (2, congruent vs. incongruent) with RT, ER and LISAS, respectively. The interaction between conflict type alternation and the CAE was our major analysis of interest.

### 2.4. Image acquisition

Functional imaging was performed on a 3T Trio scanner (Siemens Medical Systems, Erlangen, Germany) using echoplanar imaging (EPI) sensitive to blood-oxygen-level dependent (BOLD) contrast (in-plane resolution of 3.4 × 3.4 mm^2^, 64 × 64 matrix, 32 slices with a thickness of 3 mm and an interslice skip of 0.99 mm, repetition time (TR) of 2000 ms, echo-time (TE) of 30 ms, and a flip angle of 90°). In addition, a sagittal T1-weighted anatomical image was acquired as a structural reference scan, with a total of 128 slices at a thickness of 1.33 mm with no gap and an in-plane resolution of 1.0 × 1.0 mm^2^.

### 2.5. Image processing

#### 2.5.1. Preprocessing

The acquired images were processed using SPM12 software (http://www.fil.ion.ucl.ac.uk/spm/). For each subject and for each functional run, the first five volumes were discarded. The remaining images were corrected for head movement between scans by an affine registration. In one of the twenty subjects, head movements of rotation within two of four functional runs exceeded 2 degrees and therefore was excluded from further analyses. The T1 image was segmented into gray matter, white matter, cerebrospinal fluid, skin, skull and air. The head-motion-corrected functional images were aligned to the T1-weighted anatomical image through rigid-body registration. Then, the EPI images were spatially normalized to standard Montreal Neurological Institute (MNI) space using the spatial normalization parameters that mapped the structural image to the MNI space template. Normalized data were smoothed using an 8 mm full-width at half-maximum (FWHM) Gaussian kernel.

#### 2.5.2. Whole-brain analysis

For statistical analysis, fMRI data were analyzed using a two-level hierarchical general linear model (GLM). The first-level design matrix modeled fixed effects over the four sessions of smoothed data. Each session was modeled using eight event-related regressors, one for each of the conflict sequence conditions (repeated, altered, incongruent and congruent components represented by rep, alt, I and C, respectively, to define the conditions as repCC, repCI, repIC, repII, altCC, altCI, altIC, and altII). In addition, another regressor modeled errors/missed trials, and six regressors of no interest contained the realignment parameters to correct for motion artifacts. The eight conditions and the error regressors were convolved with a canonical hemodynamic response function (HRF) in SPM. Low-frequency signal drifts were filtered using a cutoff period of 128 s. Linear *t*-contrasts for CAE (CI-CC vs. II-IC) as well as the reverse contrast in conflict type repetition and alternation were tested (Chun et al., 2017; Michels, 2016). We also examined the first-order contrasts (I vs C and its reverse contrast) on average for all conditions, as well as that for type repetition and alternation conditions separately (Table 2). In the second level, one-sample t-tests of the above contrasts were analyzed. We adopted the voxel-level threshold of *p* < .005 (one-tailed) and a minimum cluster of 300 voxels (2400 mm^3^) to explore the whole-brain activities. The contrast images in volume were transferred into surface and visualized with Connectome Workbench software (Van Essen et al., 2013).

#### 2.5.3. Post hoc ROI analysis of CAE-like neural activaties

To further clarify the specific activation patterns in conflict-type repetition and alternation conditions, we performed an ROI analysis with the regions reported in the whole-brain analysis. We first tested whether each region showed a CAE activation in both conflict-type repetition and alternation conditions identified by equation (1) with one-sample *t* tests, and then extracted beta estimation values of each region (for the eight conditions) to illustrate the exact activation patterns.

To evaluate whether the neural activations of task-positive and task-negative networks could predict the corresponding behavioral performance, we took an overlap of the survival brain areas in conflict-type repetition condition and task-positive networks, including the frontoparietal network (FPN), the dorsal attention network (DAN) and the cingulo-opercular network (CON) as the task-positive areas; similarly, task negative areas were defined as the overlapping areas between the survival brain areas in conflict-type alternation condition and task-negative network (i.e., the DMN). Network atlas was adopted from Power et al. (2011).

#### 2.5.4. Connectivity analysis

The Conn toolbox (Version 19.c, Whitfield-Gabrieli & Nieto-Castanon, 2012) was used to compute the functional connectivity of different brain areas activated in different conditions. The first peak coordinates of task-positive and task-negative areas reported in the whole-brain analysis (Table 1) were selected as ROIs. The weighted GLM method was used. By convolving the HRF of the temporal BOLD signal, the ten events (eight task conditions, one error/missing condition and one rest condition) regressors and their first-order derivatives were included. In addition, six head motions as well as their first-order derivatives, the white matter and the cerebrospinal fluid were regressed out. The residuals were then used to calculate task-based functional connectivity. The connectivity values of the eight conditions of interest (i.e., repII, repIC, repCI, repCC, altII, altIC, altCI, and altCC) were averaged and then entered into second-level analysis. Standard cluster-based parametric inferences were applied to examine the clusters of functional network connectivity.

**Table 1.**
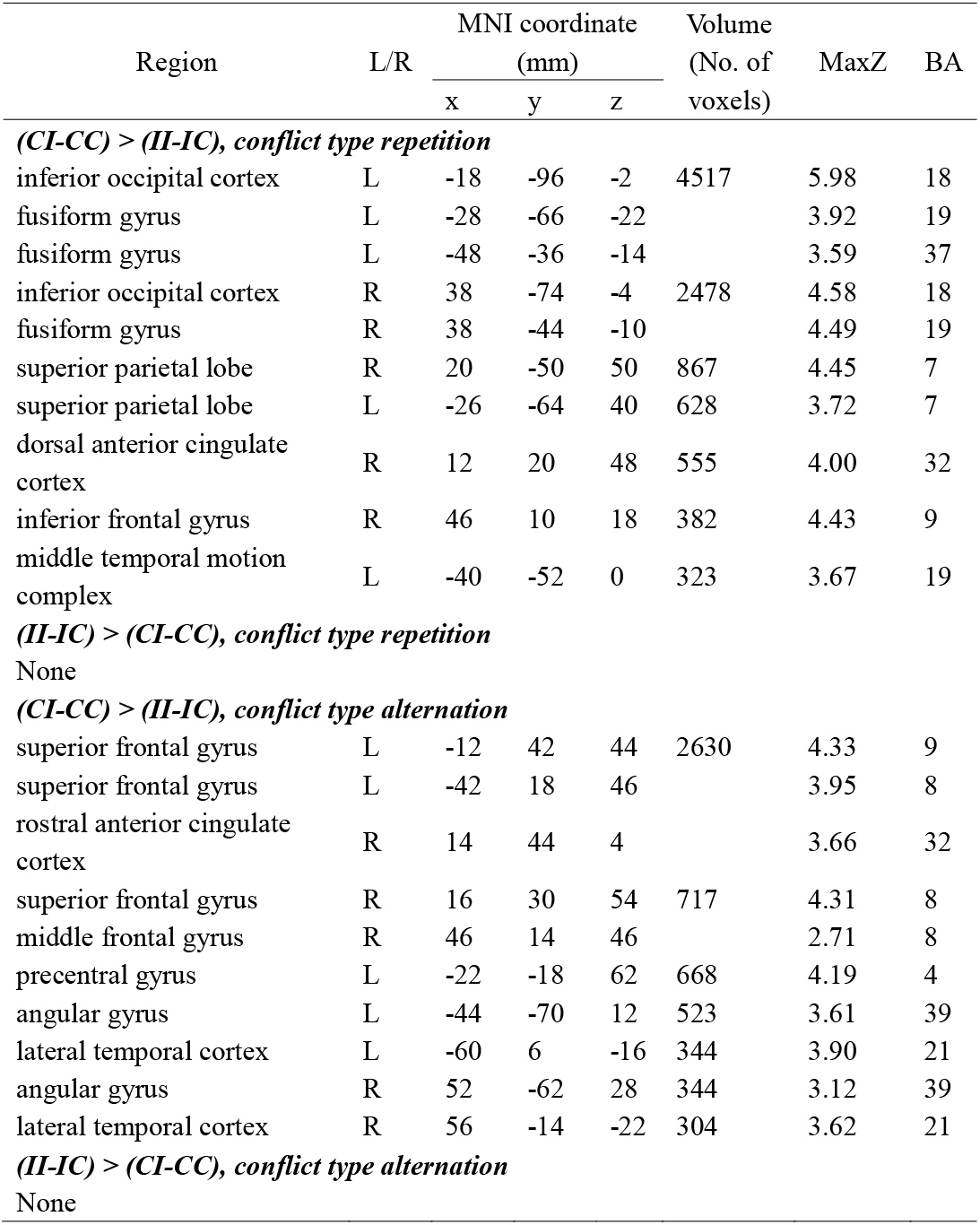
Brain activations for CAE effects in conflict type repetition and alternation conditions.

| Region | L/R | MNI coordinate |  |  | Volume<br>(No. of<br>voxels) | MaxZ | BA |
| --- | --- | --- | --- | --- | --- | --- | --- |
|  |  | (mm) |  |  |  |  |  |
|  |  | x | y | z |  |  |  |
| *(CI-CC) > (II-IC), conflict type repetition* |  |  |  |  |  |  |  |
| inferior occipital cortex | L | -18 | -96 | -2 | 4517 | 5.98 | 18 |
| fusiform gyrus | L | -28 | -66 | -22 |  | 3.92 | 19 |
| fusiform gyrus | L | -48 | -36 | -14 |  | 3.59 | 37 |
| inferior occipital cortex | R | 38 | -74 | -4 | 2478 | 4.58 | 18 |
| fusiform gyrus | R | 38 | -44 | -10 |  | 4.49 | 19 |
| superior parietal lobe | R | 20 | -50 | 50 | 867 | 4.45 | 7 |
| superior parietal lobe | L | -26 | -64 | 40 | 628 | 3.72 | 7 |
| dorsal anterior cingulate<br>cortex | R | 12 | 20 | 48 | 555 | 4.00 | 32 |
| inferior frontal gyrus | R | 46 | 10 | 18 | 382 | 4.43 | 9 |
| middle temporal motion<br>complex | L | -40 | -52 | 0 | 323 | 3.67 | 19 |
| *(II-IC) > (CI-CC), conflict type repetition* |  |  |  |  |  |  |  |
| None |  |  |  |  |  |  |  |
| *(CI-CC) > (II-IC), conflict type alternation* |  |  |  |  |  |  |  |
| superior frontal gyrus | L | -12 | 42 | 44 | 2630 | 4.33 | 9 |
| superior frontal gyrus | L | -42 | 18 | 46 |  | 3.95 | 8 |
| rostral anterior cingulate<br>cortex | R | 14 | 44 | 4 |  | 3.66 | 32 |
| superior frontal gyrus | R | 16 | 30 | 54 | 717 | 4.31 | 8 |
| middle frontal gyrus | R | 46 | 14 | 46 |  | 2.71 | 8 |
| precentral gyrus | L | -22 | -18 | 62 | 668 | 4.19 | 4 |
| angular gyrus | L | -44 | -70 | 12 | 523 | 3.61 | 39 |
| lateral temporal cortex | L | -60 | 6 | -16 | 344 | 3.90 | 21 |
| angular gyrus | R | 52 | -62 | 28 | 344 | 3.12 | 39 |
| lateral temporal cortex | R | 56 | -14 | -22 | 304 | 3.62 | 21 |
| *(II-IC) > (CI-CC), conflict type alternation* |  |  |  |  |  |  |  |
| None |  |  |  |  |  |  |  |

## 3. Results

### 3.1. Behavioral Results

For the RT, we observed a significant main effect of current congruency, *F*(1, 18) = 153.37, *p* < .001, η_*p*_^2^ = .90. Participants’ responses were slower in incongruent condition (445 ms) than in congruent condition (416 ms), indicating a conflict effect. The main effect of previous congruency was also significant, *F*(1,18) = 7.40, *p* = .014, η_*p*_^2^ = .29. Participants responded more slowly in post-incongruent conditions (432 ms) than in post-congruent conditions (429 ms), indicating a post-conflict slowing effect (Verguts, Notebaert, Kunde, & Wuhr, 2011). We also observed an interaction between previous congruency and current congruency (i.e., CAE), *F*(1,18) = 16.17, *p* = .001, η_*p*_^2^ = .47, suggesting that the conflict effect (incongruent vs. congruent) was significantly smaller after incongruent trials (445 ms vs. 413 ms) than after congruent trials (444 ms vs. 420 ms). Moreover, the interaction among consecutive conflict type, previous congruency, and current congruency was significant, *F*(1,18) =12.15, *p* = .003, η _*p*_^2^ = .40. Simple effect analyses revealed that there was a significant CAE only in the conflict type repetition condition (16 ms), *F*(1,18) = 26.19, *p* < .001, but not in the conflict type alternation condition (0 ms), *F*(1,18) < 0.01, *p* = .986. No other main effects or interactions were observed (see Figure 2A).

**Figure 2.**
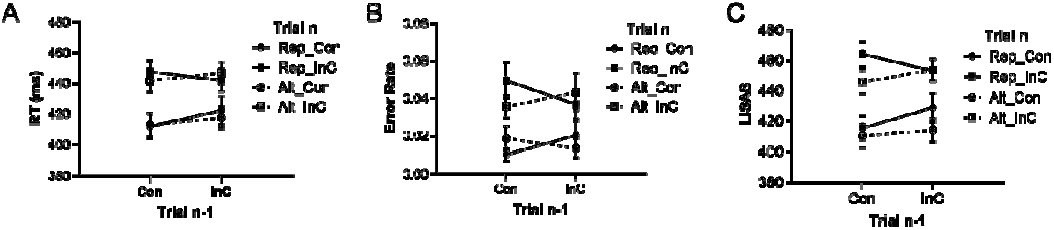
Behavioral CAE as measured by RT, ER and LISAS. When adjacent trials are of the same conflict type, CAE can be observed, i.e., an incongruent previous trial leads to a smaller conflict effect than a congruent previous trial does. In contrast, when adjacent trials are of the different conflict types, no CAE is observed. Error bars indicate standard errors. Con = congruent; InC = incongruent; Rep = repetition of conflict type; Alt = alternation of conflict type; RT = reaction time; ER = error rate; LISAS = linear integrated speed-accuracy score.

For the ER, there was a significant main effect of current congruency (i.e., conflict effect), *F*(1, 18) = 27.06, *p* < .001, η_*p*_^2^= .60. Participants had a higher ER in incongruent conditions (4.2%) than in congruent conditions (1.6%). Importantly, the interaction among consecutive conflict type, previous congruency, and current congruency was significant, *F*(1,18) =4.96, *p =* .039, η_*p*_^2^ = .22. Simple effect analyses revealed that there was a significant CAE only in the conflict type repetition condition (2.3%), *F*(1,18) = 4.91, *p* = .040, but not in the conflict type alternation condition (−1.3%), *F*(1,18) = 2.65,*p* = .121. No other significant main effects or interactions were found (see Figure 2B).

For the LISAS, there was a significant main effect of current congruency (i.e., conflict effect), *F*(1, 18) = 123.73, *p* < .001, η_*p*_^2^= .87. Participants responded more slowly in incongruent conditions (458 LISAS units) than in congruent conditions (421 LISAS units). The interaction between previous congruency and current congruency (i.e., CAE) was significant, *F*(1,18) = 13.76, *p* = .002, η_*p*_^2^= .43, suggesting that the conflict effect (incongruent vs. congruent) was smaller after incongruent trials (459 LISAS units vs. 417 LISAS units) than after congruent trials (457 LISAS units vs. 425 LISAS units). Moreover, the interaction among consecutive conflict type, previous congruency, and current congruency conditions was significant, *F*(1,18) =20.56, *p <* .001, η_*p*_^2^ = .53. Simple effect analyses revealed that there was a significant CAE only in the conflict type repetition condition (24 LISAS units), *F*(1,18) = 26.10, *p* < .001, but not in the conflict type alternation condition (−3 LISAS units), *F*(1,18) < 1. No other main effects or interactions were observed (see Figure 2C).

### 3.2. FMRI Results

#### 3.2.1. Brain activation correlates of CAEs: when conflict type repeats vs. when it changes

When the previous trial was of the same conflict type, the CAE (i.e., greater conflict effect [activation in incongruent condition minus activation in congruent condition] after a congruent trial than the conflict effect after a conflict trial) is reflected in the activation of the bilateral inferior occipital cortices (IOC), bilateral SPL, ACC, IFG, and middle temporal motion complex (MT+) (Table 1). In contrast, when the conflict type changes between consecutive trials, the conflict effect (incongruent activation minus congruent activation) after a previous congruent trial was found to be greater than that after a previous conflict trial in the bilateral superior frontal gyri (SFG), left pre-central gyrus (preCG), bilateral angular gyri (AG) and bilateral lateral temporal cortex (LTC), also showing CAE-like activities.

#### 3.2.2. Brain activation correlates of conflict effects

The average conflict effect was associated with brain areas commonly reported in conflict tasks, such as supplementary motor area, inferior parietal lobe, and so on. We also observed deactivation of superior/medial frontal regions. Further analyses showed that the activations were driven by the conflict type repetition condition, and the deactivations were driven by type alternation condition (see Table 2).

**Table 2.**
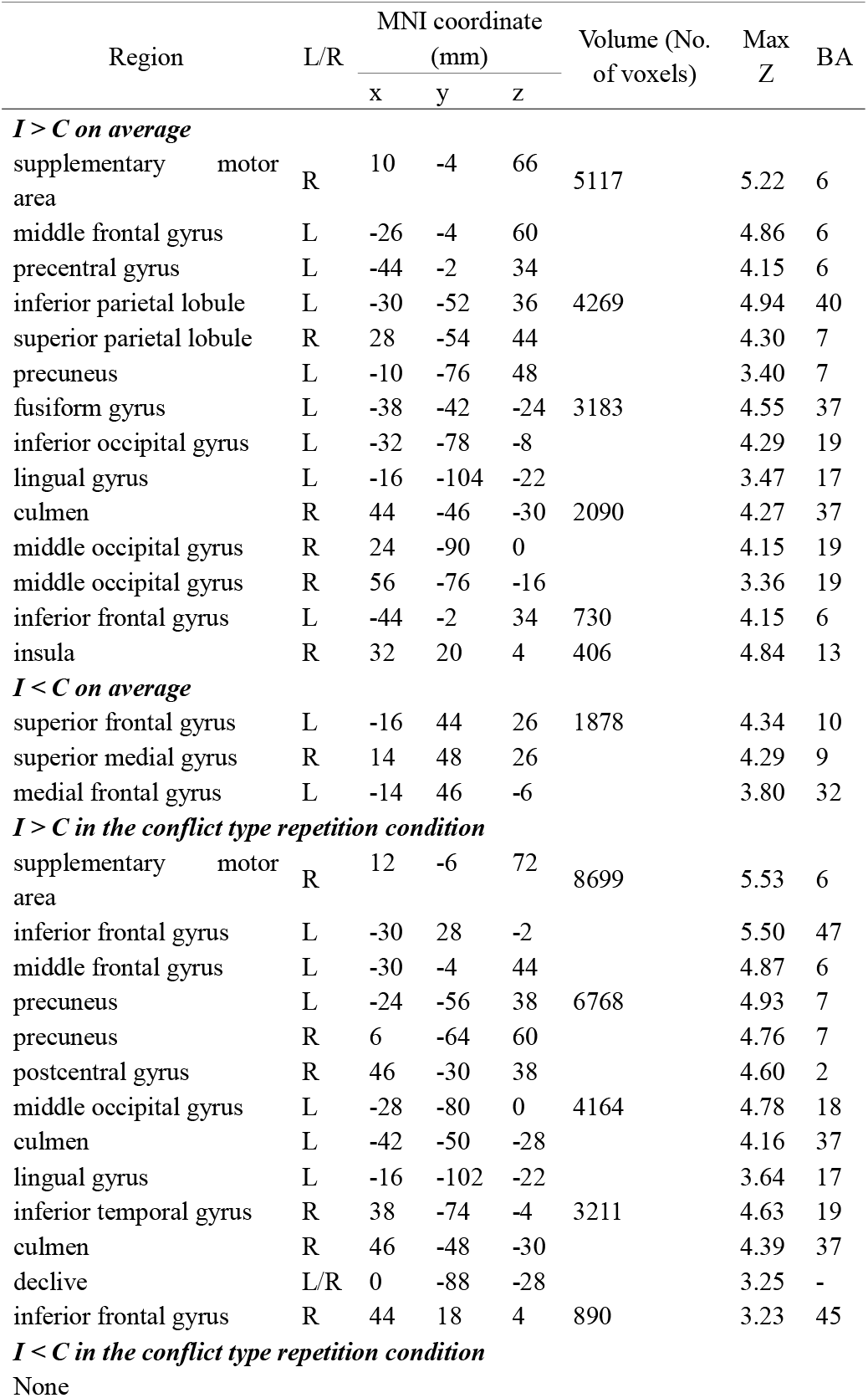
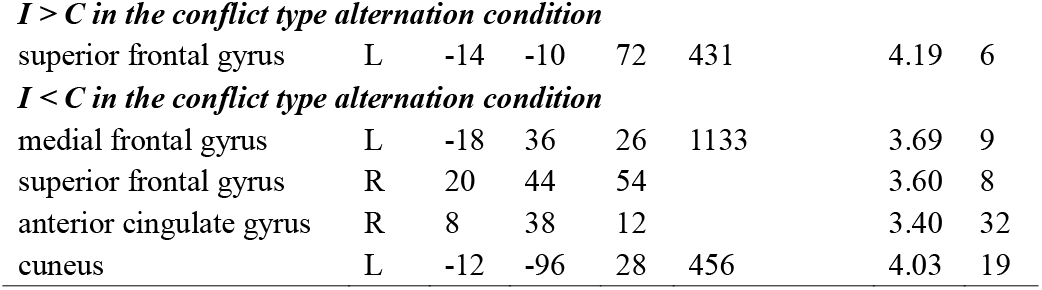
Brain activations for the first-order contrast analysis

#### 3.2.3. Post hoc ROI analysis of CAE-like neural activities

One-sample *t* test of the CAEs revealed clear dissociations between the conflict type repetition and alternation conditions (Figure 4 and 5, bar plots). On the one hand, those brain areas showing CAE-like neural activities in conflict type repetition condition were entirely inactive in conflict type alternation condition (*p*s > .110). On the other hand, those brain areas showing CAE-like neural activities in conflict type alternation condition were either inactive (for the bilateral SFG, left preCG, left AG and bilateral LTC, *p*s > .090) or deactivated (for the right AG, *p* = .032) in the conflict type repetition condition. In addition, we extracted the activations for each of the eight basic conditions (e.g., repIC, Figures 4 and 5, line graphs). We found that the areas activated in the conflict type repetition condition were positively activated, and the areas activated in the conflict type alternation condition were negatively activated in most cases.

#### 3.2.4. Verifying the involvement of task-positive/task-negative networks

In view of the above results, we investigated whether the areas showing CAE-like neural activities in the conflict type alternation condition matched the task-negative network, and the areas showing CAE-like neural activities in the conflict type repetition condition matched the task-positive networks. We applied two different methods to clarify this issue. First, the masks of task-positive and task-negative networks from Power et al.’s (2011) parcellation (see the grey areas in Figure 3A and 3B) were used to examine whether the activated areas were contained by the task-positive/negative networks. We computed the percentage of voxels inside the suspected networks, with the regions reported in Table 1, except the bilateral IOC and left preCG. For instance, we suspected that the brain areas of the ACC, IFG, MT+ and bilateral SPL activated in the conflict type repetition conditions were inside the task-positive networks. The number of voxels overlapping with the task-positive networks was 1773, and the total number of activated areas was 2755. Therefore, the brain area percentage within task-positive networks for the conflict type repetition condition was 64.4%. Similarly, the brain area percentage within the DMN for the conflict type alternation condition was 74.0% (3599/4862).

**Figure 3.**
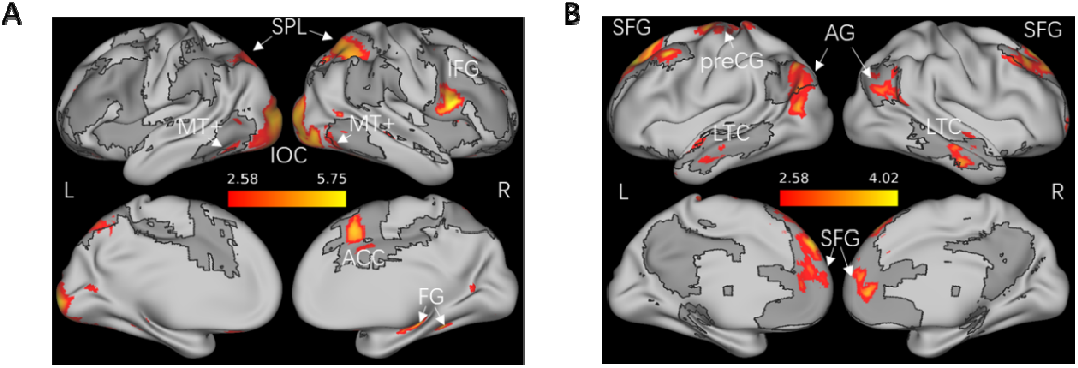
Brain correlates of the CAE for conflict-type repetition and alternation conditions respectively. Significant regions for the CAE contrast, (CI-CC)-(II-IC), are shown in A and B. The dark gray areas indicate the task-positive networks (including the dorsal attentional network, frontoparietal network and cingulo-opercular network) in (A) and task-negative network (i.e., the DMN) in (B). The templates of networks were adopted from the atlas of Power et al. (2011). Abbreviations. IOC = inferior occipital cortex; FG = fusiform gyrus; SPL = superior parietal lobe; ACC = anterior cingulate cortex, IFG = inferior frontal gyrus, MT+ = middle temporal motion complex; SFG = superior frontal gyrus; preCG = precentral gyrus; AG = angular gyrus; LTC = lateral temporal cortex; L = left; R = right.

**Figure 4.**
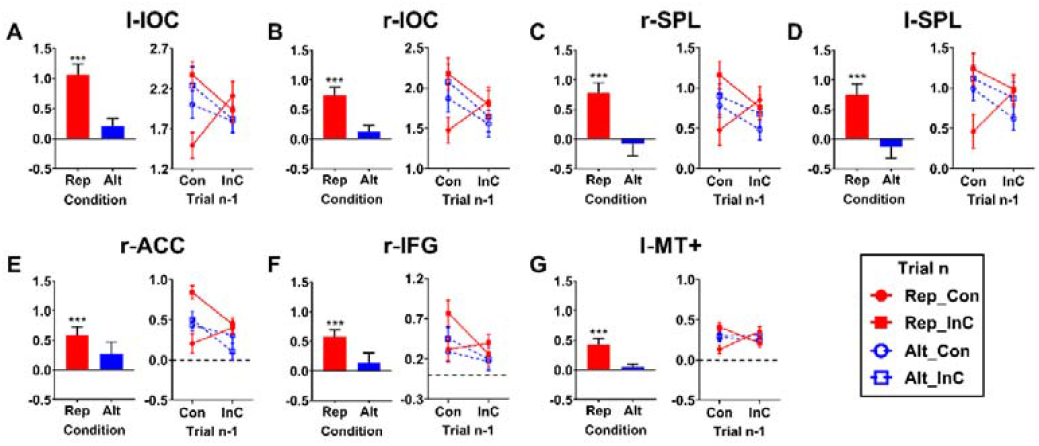
The activation of each ROI activated in conflict type repetition condition. The line graphs show the beta values as a function of congruent and incongruent conditions for both current and previous trials and their relationship (type repetition or alternation). The points above the dash lines denote positive activations. The bar plots show the CAE-like neural activities calculated by beta contrasts of (CI-CC) - (II-IC). Error bars stand for standard error. *** denotes *p* < .001; ** denotes *p* < .01; * denotes *p* < .05. Abbreviations. Con = congruent; InC = incongruent; Rep = repetition; Alt = alternation.

**Figure 5.**
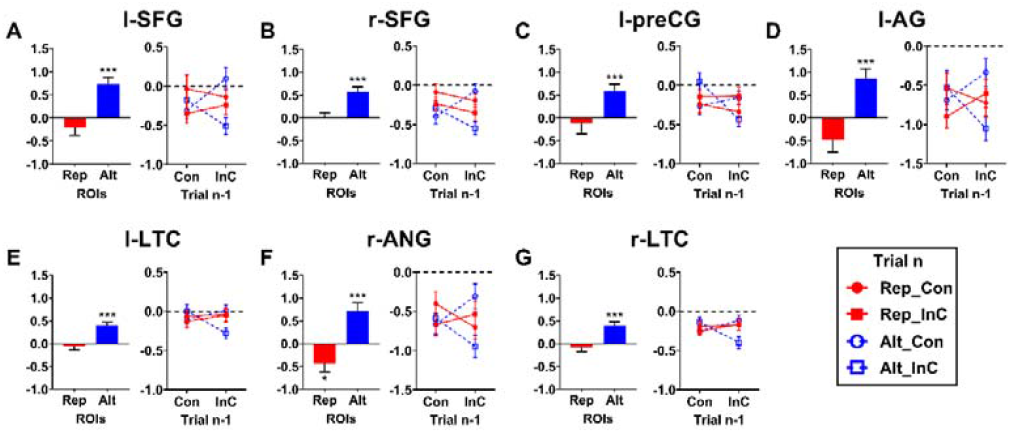
The (de)activation of for each ROI activated in conflict type alternation condition. The line graphs show the beta values as a function of congruent and incongruent conditions for both current and previous trials and their relationship (type repetition or alternation). The points below the dash lines denote negative activations (i.e., deactivations). The bar plots show the CAE-like neural activities calculated by beta contrasts of (CI-CC) - (II-IC). Error bars stand for standard error. *** denotes *p* < .001; ** denotes *p* < .01; * denotes *p* < .05. Abbreviations. Con = congruent; InC = incongruent; Rep = repetition; Alt = alternation.

To examine whether the task-positive and task-negative brain areas functioned as networks, we computed the functional connectivity between these ROIs. Connectivity analysis revealed two closely connected clusters (see Figure 6). One cluster constituted the brain areas activated in the conflict type alternation condition, namely, the bilateral AG, bilateral SFG and bilateral LTC, with 15 (i.e., a full connection, 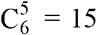) significant ROI-to-ROI connections, *F*(2,17) = 234.90, *p*-FDR = .000. The other cluster constituted the brain areas activated in the conflict type alternation condition, namely, the ACC, MT+, IFG, and bilateral SPL, with eight significant ROI-to-ROI connections (the two nonsignificant connections were IFG-MT+ and ACC-MT+), *F*(2,17) = 100.38, *p*-FDR = .000. These two clusters were significantly anti-correlated, with 29 ROI-to-ROI connections (with an exception only between r-SFG and IFG), *F*(2, 17) = 85.43, *p*-FDR = .000.

**Figure 6.**
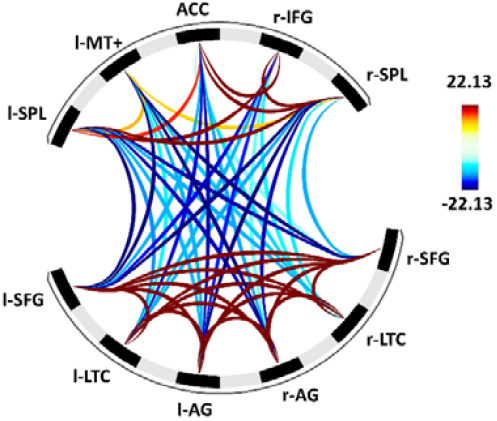
Functional connectivity of the task-positive and task-negative brain areas activated during the conflict type repetition and alternation conditions. The color denotes the t-value for each connectivity.

### 3.2.5. Correlations between brain activities and behaviors

Correlation analyses were conducted to examine whether task-positive and task-negative areas modulated the CAE size. The activated regions within task-positive and task-negative networks (by excluding the voxels outside the corresponding networks) were selected as two whole ROIs. The CAE-like neural activities of the task-positive and task-negative ROIs were calculated similarly to the behavioral CAEs (i.e., the LISAS results). We found a significant negative correlation between the task-negative ROI (de)activation and the behavioral performance in the conflict type alternation condition, *r* = -0.43, *p* = .034 (see Figure 7B). However, no correlation was observed between the average activation of task-positive areas and CAEs in the conflict type repetition condition, *r* = 0.25, *p* = .15 (see Figure 7A).

**Figure 7.**
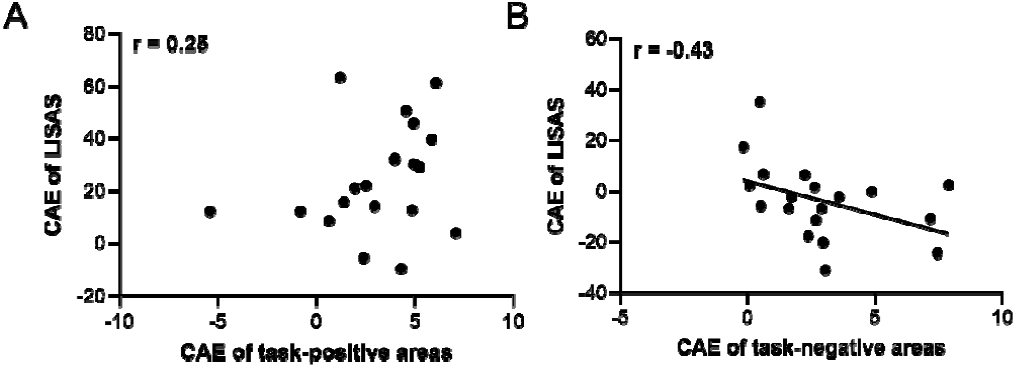
Scatter plot of the relationships between fMRI-level and behavioral-level CAE in the conflict-type repetition condition (A) and the conflict-type alternation condition (B).

## 4. Discussion

With the Stroop-color-Simon paradigm which discreetly combines the two distinct types of conflict, the present study aimed to examine the neural mechanisms underlying the sensitivity of the CAE to the coherence of conflict types. We demonstrated that with an adequate analytic strategy, CAE-like neural activities can be observed both within a conflict type and between distinct conflict types. Specifically, when the conflict type repeated (but not when it alternated), CAE-like neural activities were manifested as an activation pattern in regions within task-positive networks (i.e., the dACC, IFG, SPL and MT+). Whereas when the conflict type alternated (but not when it repeated), CAE-like neural activities were associated with a deactivation pattern in regions within task-negative networks (i.e., the SFG, AG and LTC). The CAE-like neural activities of task-negative networks could also predict the behavioral cross-type CAEs. Network analyses suggest that the two groups of brain regions showed synchronous activity within their respective group, on the other hand regions showed antagonistic activity between the two groups. To our knowledge, this is the first report on the task-negative network correlates of the sensitivity of CAEs in relation to conflict types. These findings extended our understanding of the conflict type sensitive CAE processing.

### 4.1. CAE-like neural activities in DMN Regions When Conflict Type Alternates

One novel finding of this study is that when conflict-type alternates, our neural adaptation to conflicts is related to deactivation of regions within the task-negative network, i.e., after a conflict trial of another type, these regions tend to be more de-activated in the current incongruent condition than they do in the current congruent condition.

The DMN was originally characterized as a network of regions consistently being deactivated in non-self-referential, goal-directed tasks, though later it was better known as a network that becomes active during conscious rest (Raichle, 2015). Meanwhile, many DMN regions can be activated by tasks involving certain implicit processes, such as introspection, and was considered to be the source of “mind-wandering” (Andrews-Hanna, 2012). Therefore, the deactivation of the DMN is regarded as a way to reduce internal distraction, which may act as a resource compensation mechanism in demanding tasks (Anticevic et al., 2012; Rajan et al., 2019). Considering these facts, the CAE pattern we observed that after a conflict trial of another type, DMN regions tend to deactivate further for the current conflict event, may reflects the way how our brain reacts to successive control demand of another cognitive type. As shown by the correlation results, when the control demand was larger, as indexed by the lower behavioral cross-type CAE, a stronger CAE-like neural activity in DMN (corresponding to the larger deactivation of DMN in post-incongruent condition) was observed. Therefore, the DMN might have been reactively involved in the resource compensation when conflict type alternated.

Our network analysis further suggests that activity within these DMN regions tend to be synchronous and are antagonistic to activity of the task positive network (3.2.4). It seems that the adaptive reaction of our neural system to alternating conflict events is primarily manifested as the deactivation in DMN region rather that reconfiguration in task positive regions.

### 4.2. CAE-like activities in Task-positive Regions When Conflict Type Repeats

When a conflict type repeats, the same conflict resolution mechanism is supposed to be involved. Therefore, participants needed to in real time mobilize the conflict-processing mechanism that resides in task-positive regions, causing activation in these regions which were captured by fMRI signals (M. M. Botvinick, Braver, Barch, Carter, & Cohen, 2001; Kerns et al., 2004).

The task-positive regions (i.e., dACC, IFG, SPL, MT+) we observed well replicated previous studies (Egner, 2011; Egner et al., 2007; Egner & Hirsch, 2005a; Kerns, 2006; Kerns et al., 2004; Sheth et al., 2012). The dACC is believed to play a key role in conflict detection during dynamic conflict adjustment (M. Botvinick, Nystrom, Fissell, Carter, & Cohen, 1999; M. M. Botvinick et al., 2001); the right IFG is believed to act as the source of on-line cognitive control in dynamically resolving conflicts (Egner, 2011); and the SPL and MT+ are believed to bias attention resources towards task-relevant stimuli (Egner et al., 2007; Egner & Hirsch, 2005a; Purmann & Pollmann, 2015). Moreover, we found strong intrinsic connectivity between the dACC, IFG, SPL and MT+ areas, indicating that the CAE was probably attributable to a broader conception of task-positive networks, which had been largely concealed in previous studies. This idea is consistent with a recent finding that conflict resolution involves widely distributed brain areas (Q. Li et al., 2017).

Akin to the behavioral performance, these task-positive areas showed a conflict-type sensitive feature, that is, the CAE-like neural activities were not found in these areas. These results nicely replicated previous ERP studies (Q. Li et al., 2015; Z. Li et al., 2021) that found CAE sensitivity on the conflict related N2 and P3 components, but we localized the source of domain-specific CAE with a higher spatial resolution. The inactivation of the task-positive areas in the conflict type alternation condition may provide a direct explanation for the loss of the CAE when conflict type alternated. In comparison with the previous perspectives that the dissociated cognitive control mechanisms underlying Stroop and Simon conflicts prevented the CAE from occurring (Egner, 2008; Egner et al., 2007; Egner & Hirsch, 2005b; Kim et al., 2012), we shed light on the dynamic mechanisms underlying the loss of cross-conflict CAEs.

### 4.3. Other Findings

In addition to the task-positive areas, we also observed similar conflict type sensitive activities in the visual area (i.e., IOG). This may help to resolve discrepancies regarding how cognitive control modulates sensory inputs in conflict processing. Generally speaking, conflict resolution can be achieved by either facilitating task-relevant stimuli or suppressing task-irrelevant stimuli (Z. Li, Goschl, & Yang, 2020). With a face-name Stroop task, a previous study found that the fusiform face area showed a CAE-like neural activity (similar to the results of the IOG in the conflict type repetition condition in our study) when the face was task-relevant, but not when the face was task-irrelevant (Egner & Hirsch, 2005a). Egner and Hirsch (2005a) thus proposed that the conflict resolution was achieved by facilitating task-relevant information. However, this explanation was challenged by the findings of several behavioral studies (Lee & Cho, 2013; Yang et al., 2017); these researchers observed a loss of cross-conflict CAEs when task-relevant information was kept constant while task-irrelevant information changed, which was unexpected since the repetition of task-relevant information should have produced the CAE. However, our results implied that the repetition of task-relevant information may not produce the CAE when the conflict type alternated, because the task-relevant facilitation control mechanism that supports a CAE was absent, as shown by the inactivation of task-positive and visual areas. We thus propose that the facilitation of task-relevant information does underlie the conflict processing when the conflict type repeats, but it is turned down when the conflict type alternated.

We also observed that the preCG was deactivated in the conflict type alternation condition. The preCG area is generally believed to be related to motion function. A previous study found that higher activation of the preCG contributed to a faster response (P. Wang, Fuentes, Vivas, & Chen, 2013). Moreover, decreased activity in the preCG has been related to impairments in motor preparation and execution (Spinelli et al., 2011). Therefore, the deactivation of the preCG in the post-incongruent condition in our study is probably related to post-conflict slowing, as shown in the RT results. Such a finding was consistent with a previous ERP study that localized the source of RT slowing to the precentral area (Chang, Ide, Li, Chen, & Li, 2017). Post-conflict slowing possibly reflects a speed-accuracy tradeoff to make the future response less error-prone (Weissman, 2020), and the preCG might play a key role in achieving this process.

### 4.4. Limitations

There is a notable limitation in our study. Previous studies have suggested that the CAE could be attributed to both an adjustment in top-down control and bottom-up associative learning such as feature binding (for a review, see Duthoo et al., 2014). A common practice to examine the pure cognitive control mechanisms underlying the CAE is to remove the bottom-up learning trials (e.g., Yang et al., 2017), which accounted for approximately half of the total trials in our design. To obtain greater detecting power, we did not remove these bottom-up learning trials. The basic behavioral results should not have been influenced, because there is evidence that whether the bottom-up factors were removed or not did not affect the sensitivity of the CAE to conflict type (Weissman, 2020). Although it is possible that the brain (de)activations reported in our study also reflected the processing of bottom-up learning, the observation of within-conflict CAE activations mainly in task-positive networks implied a dominant contribution of cognitive control (instead of learning). Therefore, we mainly discussed the results from the top-down control perspective. To better examine the pure cognitive control mechanisms, future studies could be designed by increasing the stimulus-response sets (Braem et al., 2014; Braem et al., 2019; Duthoo et al., 2014).

### 4.5. Conclusion

Our study found that there are different brain areas involved in the within-conflict and cross-conflict CAE. On the one hand, when conflict type repeated (rather than when it alternated), the activation of task-positive areas, such as the dACC, IFG, SPL and MT+, contributed to the within-conflict CAE. On the other hand, when the conflict type alternated (rather than when it repeated), the deactivation of task-negative areas, such as the SFG, AG and LTC, contributed to the absence of the cross-conflict CAE. These two anticorrelated networks collectively modulated the conflict type sensitive CAE.

## 5. Acknowledgement

This research was supported by the National Natural Science Foundation of China (NSFC) and the German Research Foundation (DFG) [grant numbers NSFC 62061136001 and DFG TRR-169] and China Postdoctoral Science Foundation [grant number 2019M650884].

